# Genomes of two arid-zone marsupials uncover contrasting responses to climatic change

**DOI:** 10.64898/2026.03.30.708387

**Authors:** Charles Y. Feigin, Emily Trybulec, Riley F. Ferguson, Emily L. Scicluna, Ryan Sauermann, Gabrielle A. Hartley, Rachel J. O’Neill, Andrew J. Pask

## Abstract

Small marsupials in the family Dasyuridae are a key component of Australia’s arid and semi-arid fauna, whose high species richness is proposed to reflect an opportunity-driven adaptive radiation. Despite growing interest in this group from both ecological and evolutionary perspectives, genomic data for most species is non-existent, or limited to a few marker loci. Here, we generated a chromosome-level reference genome and a *de novo* mitochondrial genome for the desert-dwelling Wongai ningaui (*Ningaui ridei*). The nuclear genome assembly is highly contiguous, with a scaffold N50 of 594.5 MB and high BUSCO gene recovery (93.8%).

Additionally, we produced a draft assembly for the related, semi-arid slender-tailed dunnart (Sminthopsis *murina*). We then used these assemblies to explore the demographic histories of these species. We find evidence for contrasting patterns of population growth during the late Pleistocene and early Holocene, corresponding with differences in local climate, potentially consistent with differences in optimal habitat. The new genomic resources and demographic findings presented here provide a foundation for future studies on adaptive specialisation in this group of Australian marsupials.

**Significance Statement:** Dasyurid marsupials are the primary carnivorous and insectivorous mammals in Australia. This family includes species such as the endangered Tasmanian devil (*Sarcophilus harrisii*) and quolls (Genus *Dasyurus*), as well as an emerging model species, the fat-tailed dunnart (*Sminthopsis crassicaudata*). Despite the species richness within dasyurids, most species remain under-studied. This is particularly true of arid and semi-arid zone species, who are often small in size, live in remote habitats and are cryptic by nature. By creating genome assemblies for two dasyurid species, this study provides resources to support phylogenetic, comparative and conservation research in arid zone marsupials. Importantly, the study’s finding that arid and semi-arid species show distinct trajectories of demographic change in response to historical climate may have implications for the resilience of locally-adapted dasyurid species to ongoing climate change.

## Introduction

Australia is one of Earth’s driest continents, with more than 70% of its landmass classified as arid or semi-arid^1^. Aridification of the continent has been a long-term process, beginning several million years ago during the mid-Miocene^2^. Yet, much of the desert landscape that dominates Australia today is thought to have arisen more recently during the late Pleistocene, with these changes shaping modern patterns of species diversity^3^. The concentration of human populations in Australia along the coasts, together with the logistical challenges of working in remote, arid regions has contributed to these habitats being relatively understudied^4^.

Small marsupial carnivores in the family Dasyuridae are a key part of Australia’s arid and semi-arid fauna. Dasyuridae, which also includes widely-known species like the Tasmanian devil, is a speciose clade, with at least 30 described arid-zone species^5^. The diversity of small dasyurids is thought to reflect a radiation in response to ecological opportunity as arid habitats have expanded across Australia^6^. These species show adaptations to such climates, including torpor and the ability to store fat in their tails^7,8^. However, there is some evidence that within clades of arid-zone dasyurids, there is relatively high niche conservatism^9^. Despite their relatively high species diversity, a notable feature of dasyurid marsupials is their exceptional degree of genomic conservation, with all examined species sharing the same chromosome complement (2N = 14), with nearly identical chromosome sizes and similar g-banding patterns^10,11^.

The subfamily Sminthopsinae is a large, speciose clade of dasyurids including dunnarts, kultarr, ningaui and planigales. These small insectivores have a wide distribution across Australia and comprise an important component of its arid-zone mammalian fauna. Taxonomy of this group is in flux, with molecular phylogenies embedding the genera *Antechynomys* and *Ningaui* (kultarr and ningaui, respectively) within the dunnart genus *Sminthopsis*, rendering the latter paraphyletic^12-14^.

Improved genomic resources for this group can benefit our understanding of their evolutionary relationships as well as how these species have been impacted by, and adapted, to the aridification of Australia. Here, we produced a chromosome-scale reference genome assembly for the desert-dwelling Wongai ningaui (*Ningaui ridei*), including both the nuclear and mitochondrial genomes. Additionally, we produced a draft assembly for the slender-tailed dunnart (*Sminthopsis murina*), a member of the dunnart sub-clade to which the genus *Ningaui* is most closely related (often called the “Murina Group”)^15^. Using our assemblies, we inferred the demographic histories of these two species, finding contrasting patterns of historical effective population size during phases of climatic change.

## Results and Discussion

### Genome assemblies

A reference genome for the Wongai ningaui (Fig. 1A) was produced using a combination of long reads and Hi-C libraries with the MaSuRCA assembler and guided by the near T2T dunnart genome assembly^16^. The nuclear assembly was approximately 3 gigabases (Gb) long, with each of the six conserved dasyurid autosomes and X chromosome represented as a single, large scaffold (Fig. 1B & C). Homology between *N. ridei* chromosomes and those of the Yellow-footed antechinus (*Antechinus flavipes*) was confirmed based on shared gene content which ranged from 97.92% on chromosome 6 to 88.55% for the X chromosome (Supplementary Table 1). Gene synteny analysis comparing the Wongai ningaui to antechinus and the eastern quoll (*Dasyurus viverrinus*), representing two Dasyurid tribes (Phascogalini and Dasyurini, respectively), showed highly conserved gene order and orientation (Fig. 1C)^17-19^. Scaffold N50 was 594.5 megabases (Mb), reflecting the large sizes of dasyurid chromosomes, and the gap percentage within scaffolds was low, 0.025%. Recovery of complete mammalian BUSCO reference genes was ∼93.8%, with only 1.6% being duplicated (Fig. 1C)^20,21^. A mitochondrial genome was assembled using Pacific Biosciences HiFi reads. The mitochondrial genome (Fig. 1D) was 17,414 basepairs (bp) in length, comparable to other marsupials^22^.

**Fig. 1.**
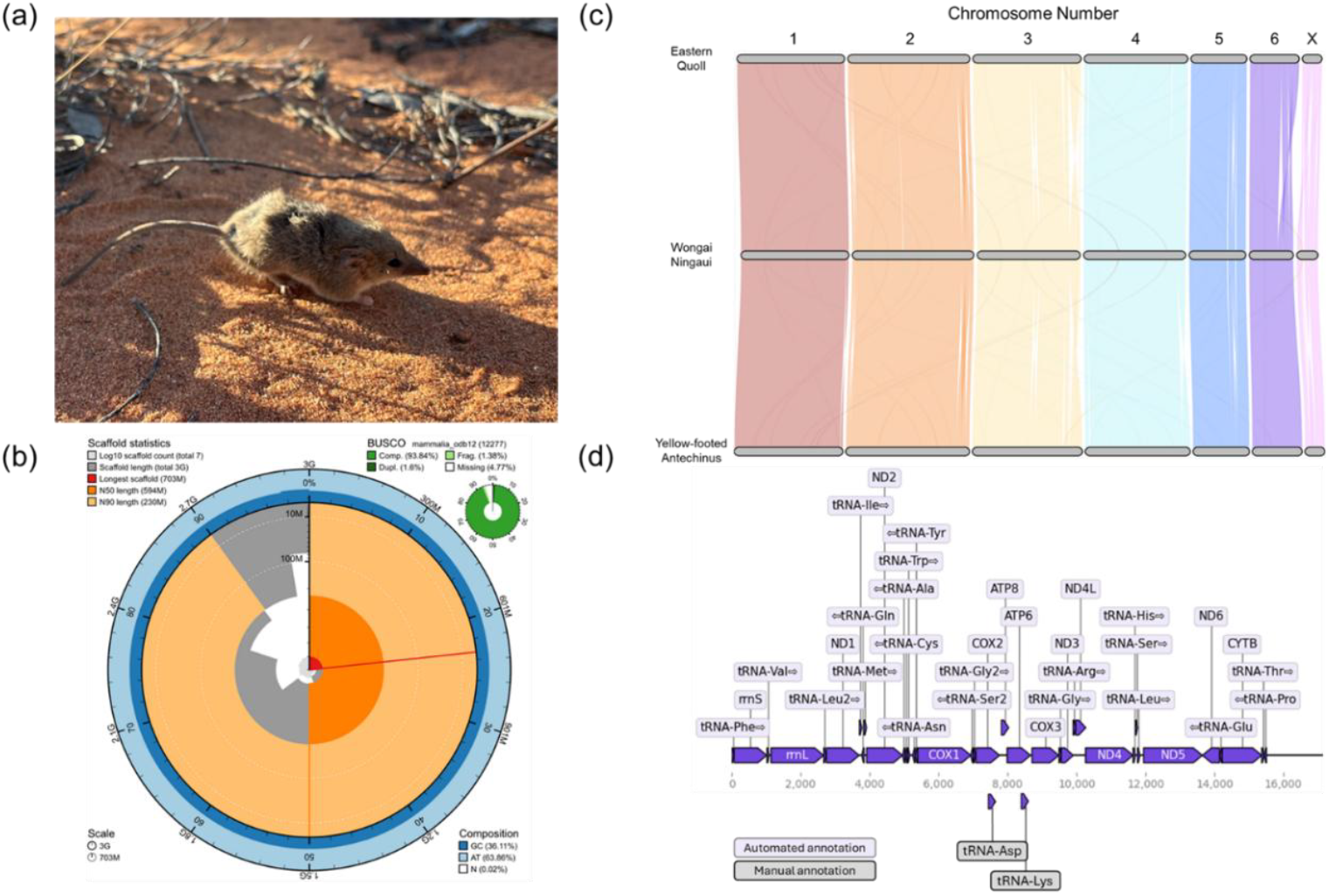
a) Photograph of a Wongai ningaui (Photo Credit: Sarah Visser). b) Snail plot displaying metrics of completeness, contiguity and base composition of the Wongai ningaui nuclear genome assembly, including scaffold statistics (count, length, N50 and N90), BUSCO gene recovery, G+C content and gaps (N content). c) Riparian plot illustrating homology and synteny between the autosomes and X chromosomes of the Wongai ningaui and the yellow-footed antechinus and eastern quoll based on homologous gene order and orientation. d) Illustration of the Wongai ningaui mitochondrial genome assembly with annotated genes.

Leveraging this reference genome, we next produced a draft nuclear assembly for the slender-tailed dunnart, based on Illumina short reads. Approximately 93.9% of contig sequence was contained within the seven chromosome-scale scaffolds (chr1-6 and the X), with the remaining 6.1% remaining in small, unplaced scaffolds (Supplementary Table 1). Given the relatively high proportion of unplaced sequences, scaffolds have a high gap percentage of 9.55%. This is comparable to that of the recent genome assembly of the extinct marsupial thylacine constructed using a similar approach^23^. BUSCO analysis recovered 90.19% of benchmarking mammal orthologs as complete and single-copy.

### Demographic histories in arid and semi-arid species

Expansion of arid zones in Australia has been associated with the retreat of many terrestrial species into refugia, particularly those with poor tolerance for such climates or with limited dispersal ability. Small dasyurids are noted both for their capacity for dispersal and their adaptations to dry environments, possibly contributing to their successful colonization across much of the arid zone^24^. However, little is known about the degree of adaptative divergence to different arid environments among small dasyurids. Understanding how populations in arid vs semi-arid environments have been affected by the progressive drying of Australia^2^ could help to predict how species may respond to continuing climate change. The Wongai and slender-tailed dunnart individuals sequenced here originate from distinctly different environments. The Wongai ningaui was collected from the central region of the Little Sandy Desert in Western Australia (WA), a true arid zone. The slender-tailed dunnart by contrast originated near Calperum Station, a mallee habitat in the Murray Darling Depression of South Australia (SA) with a semi-arid to Mediterranean climate^25^. Here, we used sequentially Markovian coalescent analysis to compare the demographic history of these species.

Our analyses suggest strikingly different trajectories of change in effective population sizes between these sister species over the late Pleistocene into the early Holocene (Fig. 2). Both species show evidence of gradually-declining Ne prior to 100,000 years ago (kya), though previous exploratory studies suggest that inferences made using this approach are less robust earlier than 100,000 generations before present (here, equivalent to 100,000 years ago assuming a 1-year generation time)^26^.

**Fig. 2.**
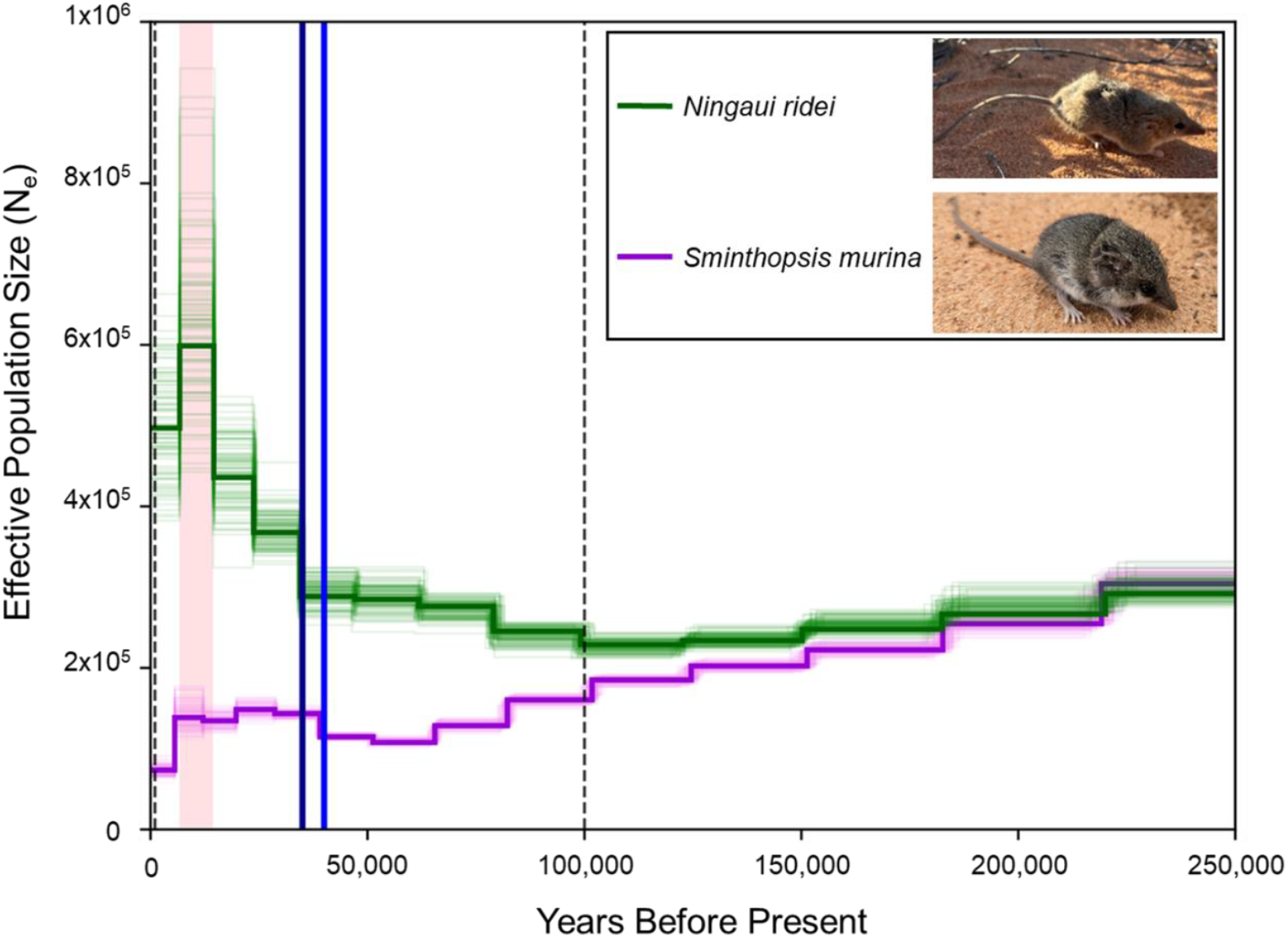
Plot showing changes in historical effective populations sizes (N_e_) in the Wongai ningaui (green) and slender-tailed dunnart (purple; photo credit Max Tibby). Thick lines represent inferences from whole genomic data, while thin lines represent bootstrap replicates. Vertical black dashed lines indicate 1,000 and 100,000 generations before present (equal to 1 kya and 100 kya), between which inferences are most reliable. Blue and dark blue vertical lines at 40 kya and 35 kya reflect the start of a humid phase and increase in fluvial activity in the Murray-Darling Basin, respectively. Pink bar indicates reducing fluvial activity 14-7kya.

Notably, from 100 kya, the two species show increasingly distinct trajectories. The Wongai ningaui showed marked increase in effective population size corresponding with gradually increasing aridity across much of Australia and accelerating during dry periods spanning much of the last 40 kya^24,27,28^. Interestingly, this pattern aligns closely with those of several other small dasyurids in partly overlapping ranges, recently inferred by extended Bayesian skyline analysis of mitochondrial data^24^. Our data may support the hypothesis that expansion of arid habitats during dry periods over the last 100 kya has promoted population growth or range expansion in arid-adapted species^24^.

In contrast to the Wongai ningaui, the slender-tailed dunnart sequenced here was inferred to have experienced almost unbroken decline from the late Pleistocene, until shortly before the Last Glacial Maximum (LGM; Fig. 2)^29^. The record of climatic change in southeastern Australia provides a less consistent picture over the last 100 kya than in northern regions, with phases of both wetter and more arid climates, though overall the region has trended toward aridity over this period. Disrupting this long-term trend, a slight demographic expansion was inferred starting just under 35 kya. This follows the onset of a humid phase in the Murray-Darling Basin around 40 kya that persisted to the LGM and increased fluvial activity at 35 kya and which ended after the resumption of drier conditions in southern Australia from ∼14-7 kya^28,30^. The contrasting trajectories of effective population size in the semi-arid slender-tailed dunnart and the desert-dwelling Wongai ningaui and their apparent correspondence to historical climatic changes may reflect evolved differences in niche preference in these species. Previous research has suggested that among small dasyurids there is modest niche conservatism^9^. While some lineages were found to have a relatively wide range of suitable climates (for example the *Sminthopsis murina* complex is found in environments ranging from mesic to semi-arid), phylogenetic clustering was highest for species in arid climates. This may reflect the intrinsic difficulty of adapting to these harsh environments. Our finding that the true arid-zone Wongai ningaui has tended to expand during periods of aridification, while *S. murina* has been negatively impacted, may support this view.

Future demographic and comparative genomic studies in these and other sminthopsine species may yield new insights into the responses of this diverse clade to accelerating climate change, as well as the biological mechanisms of, and barriers to, survival in Australia’s arid zone.

## Methods

### Tissue samples and sequencing

To sequence the Wongai ningaui genome, a small sample of muscle tissue from a juvenile female animal (Z33675) was collected in 2007 from the Little Sandy Desert of Western Australia (-24.45,122) and was acquired from the Ian Potter Australian Wildlife Biobank at Museums Victoria. DNA was extracted using the Monarch HMW DNA Extraction Kit for Tissue (NEB #T3010) per manufacturer’s instructions. DNA was precipitated from the supernatant by centrifuging the sample at 4°C for 30 minutes at 8,000 x g. The resulting pellet was washed 2x with 80% ethanol, spun at 4°C for 5 minutes at 8,000 x g in between each wash. The final pellet was eluted in nuclease-free water. DNA was quantitated using the Qubit dsDNA HS kit for the Qubit Fluorometer, and the size was determined using the Agilent Femto Pulse system. DNA was size selected using the Pacific Biosciences Short Read Eliminator (SRE) XS kit (102-208-200). For Oxford Nanopore Technologies sequencing, a library was created per the manufacturer’s instructions using the V14 sequencing kit (SQK-LSK114) with minor modifications. Modifications included extension of end repair incubations to 20 minutes at 20°C and 10 minutes at 65°C and extending all elution steps to 10 minutes at 37°C. The sample was sequenced on the ONT PromethION platform. For Pacific BioSciences HiFi sequencing, a library was created per manufacturer’s instructions using the SMRTbell prep kit 3.0 (102-141-700). The sample was sequenced on the PacBio Revio system.

Slender-tailed dunnart samples were scavenged opportunistically from a deceased, adult female animal during field work near Calperum Station, South Australia (-33.633467, 140.511497) in 2025 under Department of Energy, Environment and Climate Action permit number 10010988. Liver was collected and DNA extraction was performed using the DNEasy Blood and Tissue Kit (Qiagen). An Illumina short-insert paired-end library was produced by and sequenced to a depth of approximately 40X by the Australian Genome Research Facility using the Illumina DNA PCR-Free Assay (20041795) kit and run on an Illumina NovaSeq X Plus 25B flow cell in 2×150bp configuration.

### Genome assembly and assessment

To produce the Wongai ningaui genome assembly a total of 28.2 Gb of ONT long reads, 18.2 Gb of PacBio HiFi reads, and 130.8 Gb of Illumina short reads provided 63.3x coverage, based on the 2.8 Gb haploid genome size estimated by GenomeScope^31^. Correcting the ONT reads with Dorado resulted in reduced assembly quality. The uncorrected ONT reads, together with the PacBio and Illumina data, were used as input to the argonaut v1.0 assembly pipeline (https://github.com/emilytrybulec/argonaut). The resulting hybrid MaSuRCA assembly produced the highest completeness as indicated by Compleasm v0.2.6 and Merqury v5.2.0 score in the fewest contigs and was used moving forward^16,32,33^. To achieve chromosome-scale, the primary long read-based *de novo* assembly was then scaffolded against the closely-related fat-tailed dunnart (*Sminthopsis crassicaudata*) genome (GCA_048593235.1) using RagTag v2.1.0 without correction, leveraging the exceptional chromosomal conservation within this group^10,11,34,35^. This assembly was then polished with uncorrected ONT reads and PacBio reads using T2T autopolisher v3 to produce the 3.00 Gb final haploid assembly.

A *de novo* mitochondrial genome for the Wongai ningaui was independently assembled using MitoHiFi v3.2.1 (parameters -o 2) and the mitochondrial genome from the RefSeq Tasmanian devil mSarHar1.11 (NC_018788.1) as a reference^22^. Two genes not automatically detected by MitoHiFi (tRNA-Asp and tRNA-Lys) were annotated manually by blastn search using blast v2.17.0+ with their Tasmanian devil orthologs.

Slender-tailed dunnart short-insert, paired-end reads were pre-processed to remove residual adapters and low quality sequence using fastp v0.24.0 (parameter -- length_required 75)^36^. The assembly was generated by adapting the partial reference-guiding approach of Feigin et al. 2022^23^. First, short *de novo* contigs were assembled from paired-end reads using MEGAHIT v1.2.9 (parameter --min-contig-len 300)^37^. Duplicate contigs were removed and limited short read scaffolding was then performed using redundans v2.0.1 (paremeters --usebwa --identity 0.8 –overlap 0.8 --minLength 200 --joins 5 --limit 1.0 --iters 2)^38^. These preliminary slender-tailed dunnart scaffolds were then joined into chromosome-scale scaffolds by alignment against the SCR6 fat-tailed dunnart assembly using RagTag (parameters -f 200 -r -g 100 -m 10000000 --mm2-params ‘-x asm10’)^39^. To identify potential contaminant scaffolds, Kraken2 was used to screen the assembly, using a custom database built by adding the recently-published genome of the closely-related fat-tailed dunnart (*Sminthopsis crassicaudata*)^40^ to the “PlusPF” database, containing RefSeq archaea, bacteria, virus, vector, human, protozoa and fungi genomes^41^. This analysis identified 97.73% of sequence as marsupial, with most of the remainder unclassified (1.3%) and with only minute evidence of bacterial contamination (0.58%). Non-marsupial sequence was excluded from the final assembly. Finally, residual, unplaced mitochondrial scaffolds were identified by blastn v2.17.0+ search with default settings, using Tasmanian devil mitochondrial genome (NC_018788.1) as a query.

Contig and scaffold assembly metrics such as N50 and G+C content shown in Supplementary Table 1 were calculated using bbtools stats.sh^21^. Benchmarking Universal Single Copy Orthologs were annotated using BUSCO v6.0.0 (parameters - -mode genome --lineage mammalia_odb12). Assembly metrics for the Wongai ningaui were displayed as a snail plot using blobtoolkit v4.4.0.

### Inference of chromosome homology

Homology between ningaui chromosome-scale scaffolds and the conserved chromosome complement found across Dasyuridae was determined through comparisons of the gene content of each Wongai ningaui chromosome-scale scaffold and the chromosome scaffolds of the yellow-footed Antechinus (GCF_016432865.1 AdamAnt_v2) and eastern quoll (GCA_051122235.1 LTU_DasViv_v2.0)^17,18^. Briefly, RefSeq annotations from the yellow-footed antechinus were lifted-over to the Wongai ningaui and eastern quoll genome assemblies using LiftOff v1.6.3 with default parameters. The percentage of shared genes between each pair of approximately equal-sized chromosomes in the ningaui and antechinus genomes were calculated using perl scripts from Hartley et al. 2024^42^. Annotation overlap, along with conserved gene order and orientation between all three assemblies was then visualised as a riparian plot using GENESPACE v1.3.1. The assembly fasta files, bed files of orthologous gene locations derived from the lift-over annotations produced by LiftOff and peptide sequences extracted using gffread v 0.12.7 were provided as inputs^19,43^.

### Demographic history

Changes in effective population size of both species were inferred using the MSMC2 v 2.1.4 package^44^. Briefly, reads from each species were aligned against their respective genome assemblies using bwa-mem2 v2.2.1 (parameter -M)^45^. SAMtools v1.13 was used to process alignments, updating mate pair information (fixmate), remove duplicates (markdup, parameter -r) to filter (view, parameters -q 30 -f 3 –F 2316)^46^ and to calculate average mapping depth for each chromosome-scale scaffold (depth, parameter -a). Variants were called and mappability masks were generated using BCFtools v1.13 mpileup (parameters -q 20, -Q 20 -C 50), BCFtools call (parameters -c -V indels) and the bamCaller.py script provided with msmc-tools, providing the per-chromosome average read depths^44^. A secondary, negative mask file of repetitive regions was generated for each assembly using Red with default settings^47^. Variant calls and both mask files were provided to the generate_multihetsep.py included in msmc-tools. MSMC2 (parameters -i 50 -p 1*2+25*1+1*2+1*3) was run on the whole genome dataset and 100 bootstrap samples generated using the msmc-tools multihetsep_bootstrap.py script (parameters --seed 2025 -n 100). Data was plotted using a per-generation mutation rate of 5.95e^-9^, based on parent-offspring trio data from the closest available relative, (the Tasmanian devil) and a generation time of 1 year^48-50^.

## Author Contributions

C.Y.F., A.J.P and R.O. conceived the study. G.A.H. performed DNA extractions and sequencing of Wongai ningaui samples. E.T. and C.Y.F. assembled the Wongai ningaui and slender-tailed dunnart nuclear genomes, respectively. C.Y.F. assembled the Wongai ningaui mitochondrial genome. C.Y.F. and E.T. performed assembly assessment. C.Y.F. performed homology analyses. E.L.S. and R.S. conducted field work and acquired slender-tailed dunnart samples. C.Y.F. and R.F.F. performed demographic history analyses. C.Y.F. wrote the manuscript with contributions and input from all authors.

## Acknowledgements

We thank Kevin Rowe, Joanna Sumner and Karen Roberts and Museums Victoria for providing tissue from the Wongai ningaui, Patricia A. Woolley who collected the sample, Australian Landscape Trust who manage Calperum Station for allowing us to use the slender-tailed dunnart specimen, Sarah Visser and Max Tibby for permission to use their photos of the Wongai ningaui and slender-tailed dunnart, respectively, the Center for Genome Innovation and Computational Biology Core in the Institute for Systems Genomics at the University of Connecticut for sequencing and computational support, respectively, Nicole Pauloski for troubleshooting ONT runs and Elise Ireland for extensive proofreading. We thank the various Traditional Owners of the many lands in which these species live and that these specimens originated from.

## Data Accessibility

Wongai ningaui and slender-tailed dunnart data are available on NCBI under PRJNA1434504 and PRJNA1431567, respectively. Gene annotations and code generated in this study are available at https://github.com/charlesfeigin/Arid-zone-dasyurids-demography-paper-repo.

## AI Usage Statement

No artificial intelligence was used in the writing of this manuscript. GPT-5 was used to refactor an existing python script from Hartley et al. 2024 for plotting MSMC2 results (plot_msmc2_bootstrap.py) to improve readability.

## Funding

This study was supported through philanthropic support by the Wilson Family Trust and industry funding from Colossal Biosciences Inc to A.J.P. and by the Colossal Foundation to R.J.O.

## Conflict of Interest Statement

The authors do not declare any conflicts of interest.

## Notes

https://github.com/charlesfeigin/Arid-zone-dasyurids-demography-paper-repo

### Competing Interest Statement

The authors have declared no competing interest.

### Summary of Updates

1) Minor spelling/grammatical issues were fixed 2) A line indicating the corresponding author was added. 3) NCBI submission numbers were updated with NCBI BioProject IDs. 4) Manuscript was converted to double-spacing with line numbers added 4) Figure 2 caption was shortened by removing redundant information with main text.

